# SRF is a non-histone methylation target of KDM2B and SET7 in the regulation of myogenesis

**DOI:** 10.1101/2020.04.17.046342

**Authors:** Hosouk Joung, Joo-Young Kang, Ji-Young Kim, Duk-Hwa Kwon, Anna Jeong, Hyun-Ki Min, Sera Shin, Yun-Gyeong Lee, Young-Kook Kim, Sang-Beom Seo, Hyun Kook

## Abstract

Demethylation of histone lysines, one of the most important modifications in transcriptional regulation, is associated with various physiological states. KDM2B is a histone H3K4, H3K36, and H3K79 demethylase associated with the repression of transcription. Here, we present a novel mechanism by which KDM2B demethylates serum response factor (SRF) K165 to negatively regulate muscle differentiation, which is counteracted by histone methyltransferase SET7. We show that KDM2B inhibited skeletal muscle differentiation by inhibiting the transcription of SRF-dependent genes. Both KDM2B and SET7 regulated the balance of SRF K165 methylation. SRF K165 methylation was required for the transcriptional activation of SRF and for the promoter occupancy of SRF-dependent genes. SET7 inhibitors blocked muscle cell differentiation. Taken together, these data indicate that SRF is a non-histone target of KDM2B and that the methylation balance of SRF maintained by KDM2B and SET7 plays an important role in muscle cell differentiation.

## Introduction

Proteins are susceptible to diverse covalent modifications of their amino acid residues. These post-translational modifications (PTMs) are involved in normal homeostasis and in pathologic conditions such as cancer (Audia & Campbell, 2016; Chervona & Costa, 2012). Protein methylation, the addition of a methyl group to target proteins, is an important PTM and plays important roles in various human pathophysiology (Han *et al*, 2019; Rowe *et al*, 2019; Wei *et al*, 2014). Protein methylation is well balanced by two sets of enzymes: methyltransferases and demethylases. Histone methylation occurs on the nitrogen-containing side chains of arginine or lysine. In contrast to arginine methylation, which is mediated by a series of proteins called PRDMs that contain the PR (PRD1-BF1 and RIZ homology) domain, lysine is methylated by a family of methyltransferases with a SET [Su(var)3-9, Enhancer-of-zeste and Trithorax] domain (Schneider *et al*, 2002; Yang & Bedford, 2013).

Serum response factor (SRF) is critical in cellular survival and differentiation (Chervona & Costa, 2012; Croissant *et al*, 1996; Han *et al*., 2019; Kim *et al*, 2009). SRF directly binds to the serum response element (SRE) in the promoter of its target genes (Croissant *et al*., 1996; Kim *et al*., 2009; Tamura *et al*, 2018). Many of the biological activities of SRF, such as DNA-binding, dimerization, and interaction with other transcription factors, take place in the conserved MADS domain (Norman *et al*, 1988; Pellegrini *et al*, 1995). The MADS-box is a motif in many DNA-binding transcription factors and the sequences are well-conserved in eukaryotic organisms from yeasts to mammals. The name refers to the 4 originally identified members: MDM1, AG, DEFA, and SRF (Shore & Sharrocks, 1995). In mammals, SRF participates in the transcription of c-*fos*, a protooncogene, and initiates cell-type specific genes. In skeletal muscle cells, SRF works as a transcription factor to induce SRF-dependent muscle-specific genes (Croissant *et al*., 1996; Kim *et al*., 2009; Shore & Sharrocks, 1995). In addition to DNA-binding, the MADS-box recruits other transcription factors to form multi-component regulatory complexes. We also found that enhancer of polycomb, a chromatin protein, directly binds to SRF to potentiate the activation of MRF (Kim *et al*., 2009).

Among the methyltransferases, SET7 (SET Domain Containing 7) is a well-known H3K4 monomethyltransferase (Xiao *et al*, 2003). In addition to histones, SET7 also methylates non-histone proteins by binding to the consensus recognition motif (K/R-S/T/A-K) in substrate proteins. For example, SET7 monomethylates K189 of TAF10 and increases its affinity for RNA polymerase II (Francis *et al*, 2005; Kouskouti *et al*, 2004). It also methylates K372 of p53/TP53 to stabilize the protein (Chuikov *et al*, 2004). In addition, SET7 interacts with MyoD and SRF when myoblasts differentiate into myotubes, which is required for skeletal muscle development (Tao *et al*, 2011; Tuano *et al*, 2016).

KDM2B, also known as JHDM1B/FBXL10, is an Fe(II)- and alpha-ketoglutarate-dependent histone demethylase. It consists of multiple functional domains: an N-terminal JmjC domain, a CXXC zinc finger domain, a PHD domain, an F-box domain, and seven leucine-rich repeats. The JmjC domain catalyzes the demethylation of H3K4me3 and H3K36me2, leading to the transcriptional repression of target genes (He *et al*, 2008; Janzer *et al*, 2012). Recently, we also discovered that KDM2B demethylates H3K79 in a sirtuin-1-dependent manner (Kang *et al*, 2018). KDM2B participates in many aspects of normal cellular processes, such as cell senescence, cell differentiation, and stem cell self-renewal. Recent studies have also shown that KDM2B is overexpressed in various types of cancers (Yan *et al*, 2018; Zheng *et al*, 2018). KDM2B promotes cancer cell proliferation, metastasis, cancer stem cell self-renewal, and drug resistance (Yan *et al*., 2018; Zheng *et al*., 2018). However, KDM2B can also decrease cancer cell proliferation by inhibiting the expression of oncogenes (Yan *et al*., 2018). Other than regulating cell proliferation and thereby tumor biology, KDM2B inhibits adipogenesis as part of polycomb repressive complex 1 (PRC1), but in a demethylase-independent manner (Inagaki *et al*, 2015). However, the role of KDM2B histone demethylase in myogenic differentiation remains unknown.

In the current work, we report a novel mechanism of KDM2B-mediated demethylation of serum response factor (SRF) K165, which counteracts SET7-induced methylation. The methylation of SRF is required for the transcriptional activation of muscle-specific genes by alteration of its binding affinity to the *serum response element* (*SRE*) in the promoters of its target genes.

## Results

### KDM2B is expressed in the early phase of skeletal muscle differentiation

We first examined the distribution of KDM2B in the tissues of adult mice. KDM2B was highly expressed in the brain (Fig EV1A) as previously reported (Inagaki *et al*., 2015). To investigate the role of KDM2B in skeletal muscle, we first checked its expression using a C2C12 myoblast cell line. As shown by western blot analysis, differentiation media (DM) reduced the expression of KDM2B in C2C12 cells (Fig 1A). *Kdm2b* expression was also reduced as evidenced by quantitative RT-PCR (Fig 1B). Mouse skeletal muscles were harvested at different ages and used for western blot analysis. KDM2B was highly expressed at embryonic day 18 (E18), and expression was maintained for 1 month after birth, after which further expression gradually decreased with age (Fig 1C).

**Figure 1.**
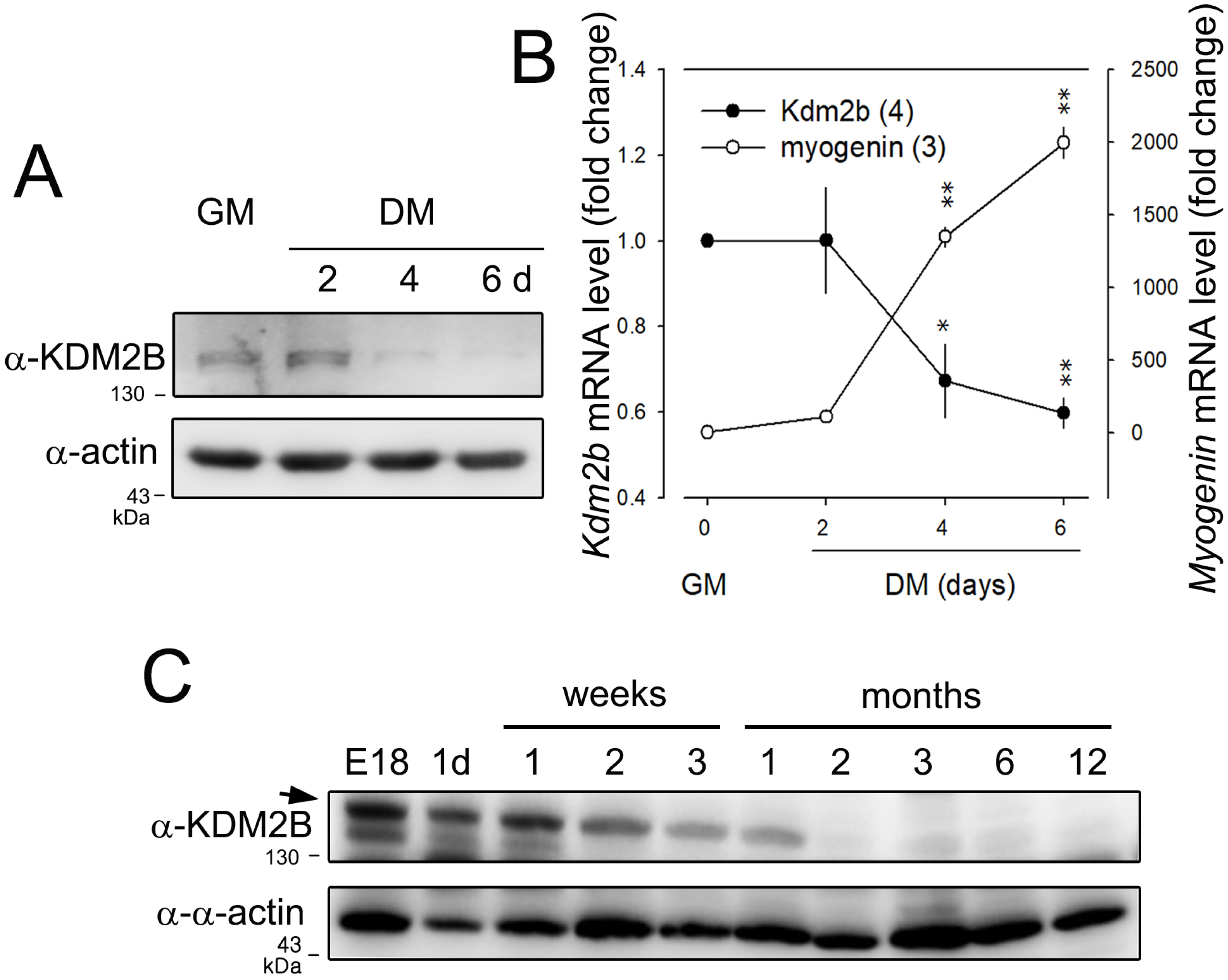
*Kdm2b* is expressed in the early phase of skeletal muscle development. A. KDM2B expression is gradually reduced in differentiation media (DM) in a C2C12 myoblast cell line. GM: growth media. B. Quantitative real-time PCR (qRT-PCR) showing that *Kdm2b* expression is down-regulated 4 and 6 days after treatment with DM in C2C12 cells. By contrast, *Myogenin*, a myogenic transcription factor, is increased during myoblast differentiation. Numbers in parentheses are the number of independent observations. *p < 0.05, **p < 0.01 compared with mRNA levels in GM. C. Mouse tissue blot analysis showing that KDM2B is abundant in the early phase of skeletal muscle maturation. Note that KDM2B is gradually down-regulated after birth.

### KDM2B inhibits skeletal muscle cell differentiation

We next examined the effect of KDM2B in C2C12 cells. Replacement of growth media (GM) with DM for 3 days caused an increase in the expression of skeletal proteins, such as myosin heavy chain (Mhc) and muscle creatinine kinase (Mck). Those increases were blunted when KDM2B was overexpressed (Fig 2A). mRNA transcript levels of skeletal *α*-actin (actin alpha 1 skeletal muscle, *Acta1*) and *Mck*, both of which are the terminal differentiation markers in myogenesis, were downregulated by transfection of *KDM2B* (Fig 2B). Expression of MyoD, a key myogenic transcription factor (Berkes & Tapscott, 2005; Yamamoto *et al*, 2018), was also significantly lowered by KDM2B (Fig 2A), as was that of *Myogenin* (Fig 2B). Treatment with DM for 3 days induced elongation (Fig 2C) and multinucleation of C2C12 cells, a key feature of myogenesis, which was attenuated by transfection of *KDM2B* (Fig 2D).

**Figure 2.**
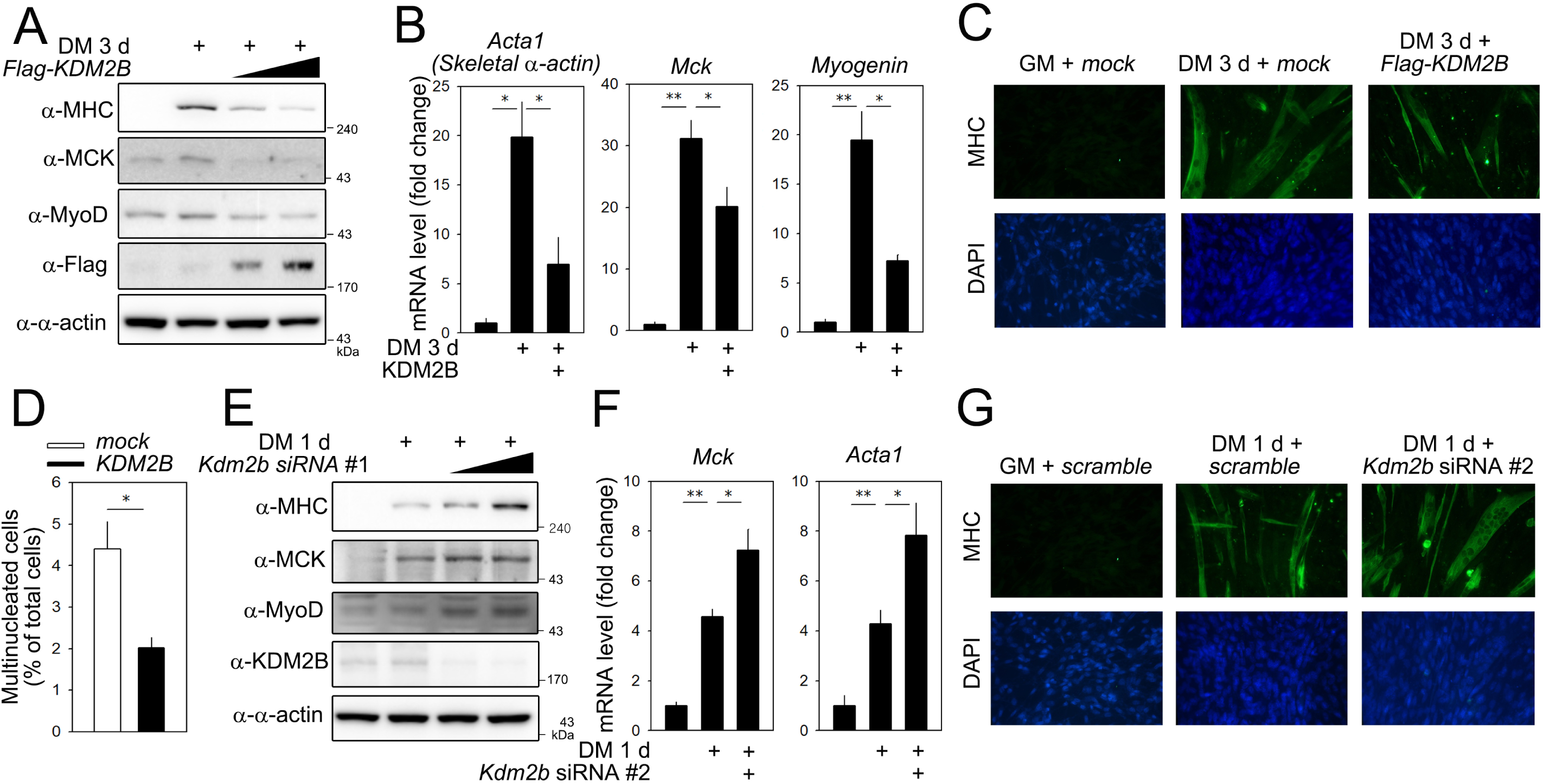
KDM2B inhibits myoblast differentiation. A. Forced expression of KDM2B decreases expression levels of skeletal muscle genes in C2C12 cells. Treatment with DM for 3 days induces expression of myosin heavy chain (Mhc), MyoD, and muscle creatinine kinase (Mck). These increases are attenuated when *Flag-tagged KDM2B* is transfected. B. qRT-PCR analysis showing that the increased transcript levels of *skeletal α-actin (Acta1), Mck*, and *Myogenin* are down-regulated by overexpression of KDM2B. C. Treatment with DM for 3 d induces elongation and multinucleation of C2C12 cells, which is attenuated by forced expression of KDM2B. Immunocytochemistry was performed with anti-MHC antibody. D. Direct cell count after immunocytochemical analysis with MHC showing that multinucleated cell count is reduced when *KDM2B* is transfected. Images from 20∼24 different fields were randomly obtained and multinucleated cells and total cells were counted. E. Knock-down of *Kdm2b* with *Kdm2b* siRNA #1 enhances the skeletal muscle gene expression induced by treatment with DM for 1 d. (F) qRT-PCR results for *Mck* and *Acta1*. G. Immunocytochemical analysis with anti-MHC antibody showing that elongation and multinucleation are enhanced by transfection of *Kdm2b* siRNA. *p < 0.05, **p < 0.01.

We next examined the effect of knocking-down *Kdm2b* using siRNA. We first introduced *Kdm2b* siRNA after treatment with DM for 3 days. However, no clear changes were observed (data not shown), which may suggest the full activation of skeletal muscle genes. Thus, we tried to check the effect of *Kdm2b* siRNA during the early phase of muscle cell differentiation. DM for 1 day induced expression of MHC and this was enhanced by *Kdm2b* siRNA (Fig 2E, *Kdm2b* siRNA #1). The increases in MCK and MyoD expressions were also potentiated (Fig 2E). To rule out an off-target effect, we designed another siRNA targeting a different location (*Kdm2b* siRNA #2). This siRNA also significantly induced the expression of myogenic proteins (Fig EV1B). The increase in the mRNA levels of *Mck* and *Acta1* were enhanced by *Kdm2b* siRNA (Fig 2F). Immunocytochemical analysis further showed that the *Kdm2b* siRNA increased the formation of skeletal myofibers in C2C12 cells induced by 1 day of DM (Fig 2G).

### KDM2B reduces transcriptional activity of myogenic genes by detaching SRF from the *SRE*

To examine the effect of KDM2B on the promoter activity of myogenic genes, we utilized promoter luciferase constructs and transfected the mammalian expression vector of KDM2B into C2C12 cells. Transfection of *Kdm2b* decreased the basal promoter activities of *Acta1* and *Mck* (Fig 3A and 3B) in a dose-dependent fashion. *Kdm2b* also attenuated the basal promoter activity of *Myogenin* (Fig EV2A). We also checked whether KDM2B affected SRF-induced transactivation (Belaguli *et al*, 1997; Fluck *et al*, 2000). SRF-induced transactivation of *Acta1* (Fig 3C), *Mck* (Fig 3D), and *Myogenin* (Fig EV2B) promoters was significantly attenuated by the transfection of *KDM2B*. Chromatin immunoprecipitation (ChIP) analysis showed binding of endogenous SRF to either the proximal or distal SRE in the *Acta1* promoter region was reduced by transfection of *KDM2B* in C2C12 cells cultured in DM for 1 day (Fig 3E and Fig EV3A). Likewise, the binding of exogenous HA-tagged SRF to *SRE* was reduced by transfection of *KDM2B* in C2C12 cells cultured in DM for 1 day (Fig EV3B and EV3C). In contrast, knocking down *Kdm2b* in C2C12 cells enhanced the binding of SRF to either the proximal or distal *SRE* in the *Acta1* promoter (Fig EV3D).

**Figure 3.**
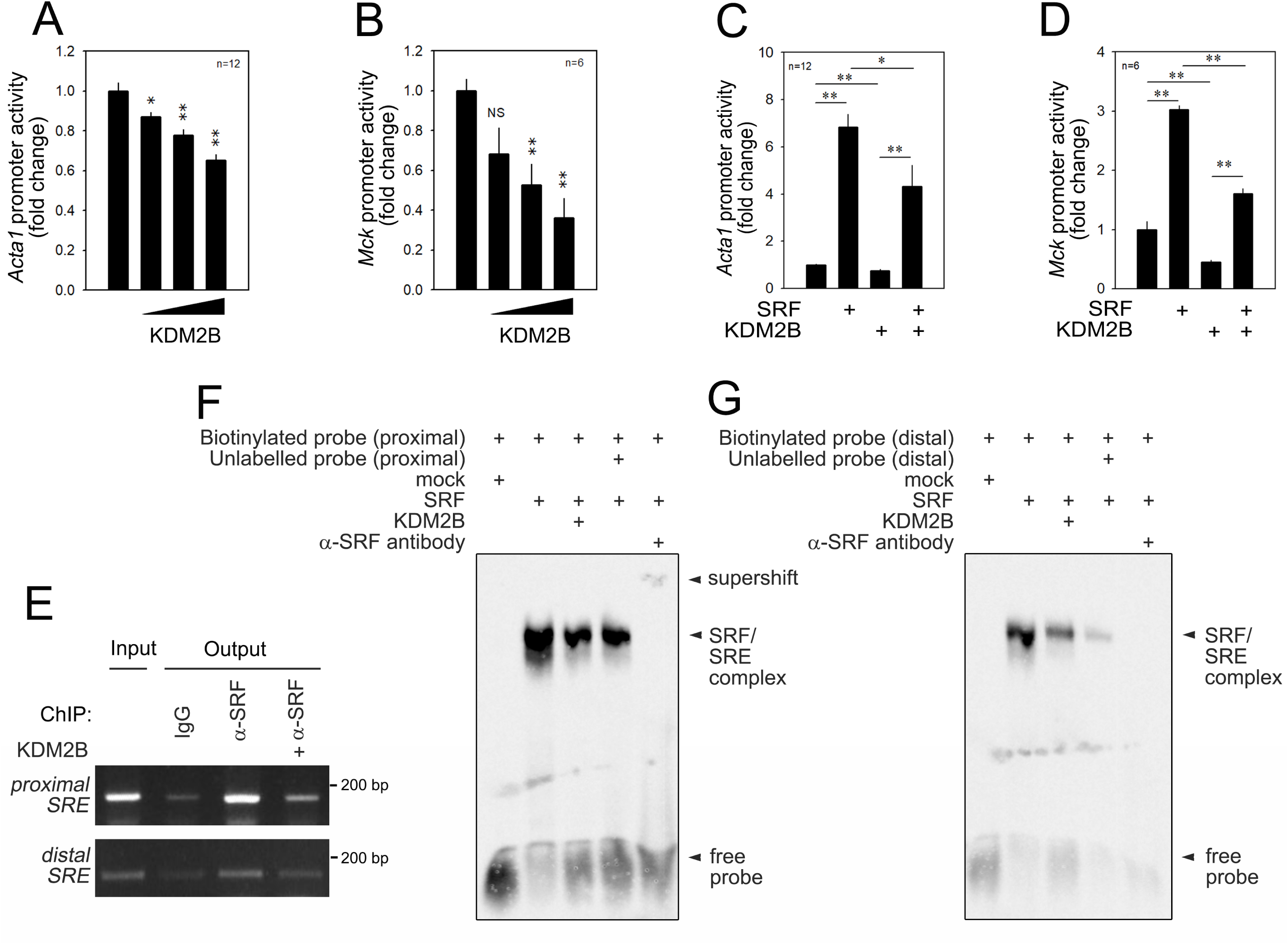
KDM2B transcriptionally inhibits serum response factor (SRF)-dependent muscle gene expression by detaching SRF from the *SRE*. A,B. Transfection of *KDM2B* reduces the basal promoter activities of *Acta1* (A) and *Mck* (B) in a dose-dependent fashion. C,D. SRF-induced transactivation of *Acta1* (C) and *Mck* (D) promoters is significantly attenuated by co-transfection with *KDM2B*. E. Chromatin immunoprecipitation (ChIP) showing that KDM2B overexpression reduces SRF occupancy at the proximal or distal serum response element (*SRE*) of the *Acta1* promoter, indicating that KDM2B induced detachment of SRF from the *SRE*. F. Gel shift assay showing that binding of SRF to the proximal *SRE* was attenuated by addition of KDM2B. Nuclear extracts of C2C12 cells transfected with *HA-SRF* and/or *Flag-KDM2B* were used. G. Gel shift assay with distal *SRE*. *p < 0.05, **p < 0.01. NS, not significant.

We also performed a gel shift assay with biotinylated *SRE* probes. Nuclear extracts from *SRF*-transfected C2C12 cells successfully retarded the gel mobility of the proximal *SRE* probe, which forms an SRF-SRE complex (2^nd^ lane, Fig 3F, arrowhead). However, the binding of SRF to the *SRE* was downregulated when nuclear extracts from the cells co-transfected with *KDM2B* and *SRF* were added (3^rd^ lane). Un-biotinylated cold probe reduced the SRF-SRE complex (4^th^ lane). Addition of *α*-SRF antibody significantly regarded the gel shift, which formed a supershifted band (5^th^ lane, arrowhead). When the distal *SRE* probe was used, we observed the same result that KDM2B reduced the formation of the SRF-SRE complex (Fig 3G). These results suggested that the transcriptional repression induced by KDM2B is caused by KDM2B-induced detachment of SRF from the *SRE*.

### KDM2B inhibits transcription in a histone-methylation-independent manner

Many researchers, including us, have demonstrated that KDM2B primarily targets histones to demethylate lysine residues (He *et al*, 2011; Kang *et al*., 2018; Yan *et al*., 2018) and that histones H3K4me3 and H3K36me2 in particular can be effectively demethylated (He *et al*., 2008; Janzer *et al*., 2012). This raises the possibility that our results showing the inhibition of myogenic transcription factors may result from KDM2B-mediated repressive modification by demethylation at H3K4 and H3K36. Thus, using ChIP analysis, we checked the methylation status of histones associated with the *SRE* in the promoter of *Acta1*. The histone H3K4me3 level associated with the proximal *SRE* in the *Acta1* promoter was not altered by transfection of KDM2B in C2C12 cells exposed to DM for 1 d. Likewise, KDM2B also did not affect the binding of H3K4me3 to the distal *SRE* (Fig EV4A and EV4B). Likewise, H3K36me2 associated with either the proximal or the distal *SRE* was not significantly altered by KDM2B overexpression (Fig EV4C and EV4D). We also found that the knocking-down of *Kdm2b* with siRNA did not increase the ‘active’ methylation of histones H3K4me3 (Fig EV4E and EV4F) or H3K36me2 (Fig EV4G and EV4H) in the two *SRE*s of *Acta1* in C2C12 cells in the GM condition. These results suggested that histone methylation is not involved in the KDM2B-mediated inhibition of transcription.

### KDM2B directly binds to myogenic transcription factors

In our experimental model, KDM2B-mediated transcriptional repression of myogenic genes is not caused by regulation of histone methylation. Many epigenetic regulators can induce modifications of proteins, including transcription factors other than histones. Thus, we were also interested in whether KDM2B directly affects the post-translational modification of key transcription factors in myogenesis. SRF is ubiquitous and has diverse functions in cell survival and differentiation; it is also critical in the early phase of skeletal muscle cell differentiation because it is required for the transcriptional activation of muscle-specific genes (Croissant *et al*., 1996) that contain a well-conserved CArG box binding motif [CC(A/T)6GG] in their promoter regions (Hauschka, 2001). Thus, we investigated whether KDM2B affects the PTM of SRF and thereby its transcriptional activity. In C2C12 myoblast cells, endogenous KDM2B physically interacted with endogenous SRF (Fig 4A). We also examined the interaction with exogenous proteins of KDM2B and SRF. Flag-tagged KDM2B successfully recruited HA-tagged SRF (Fig 4B). Likewise, HA-tagged SRF pulled down Flag-tagged KDM2B (Fig 4C).

**Figure 4.**
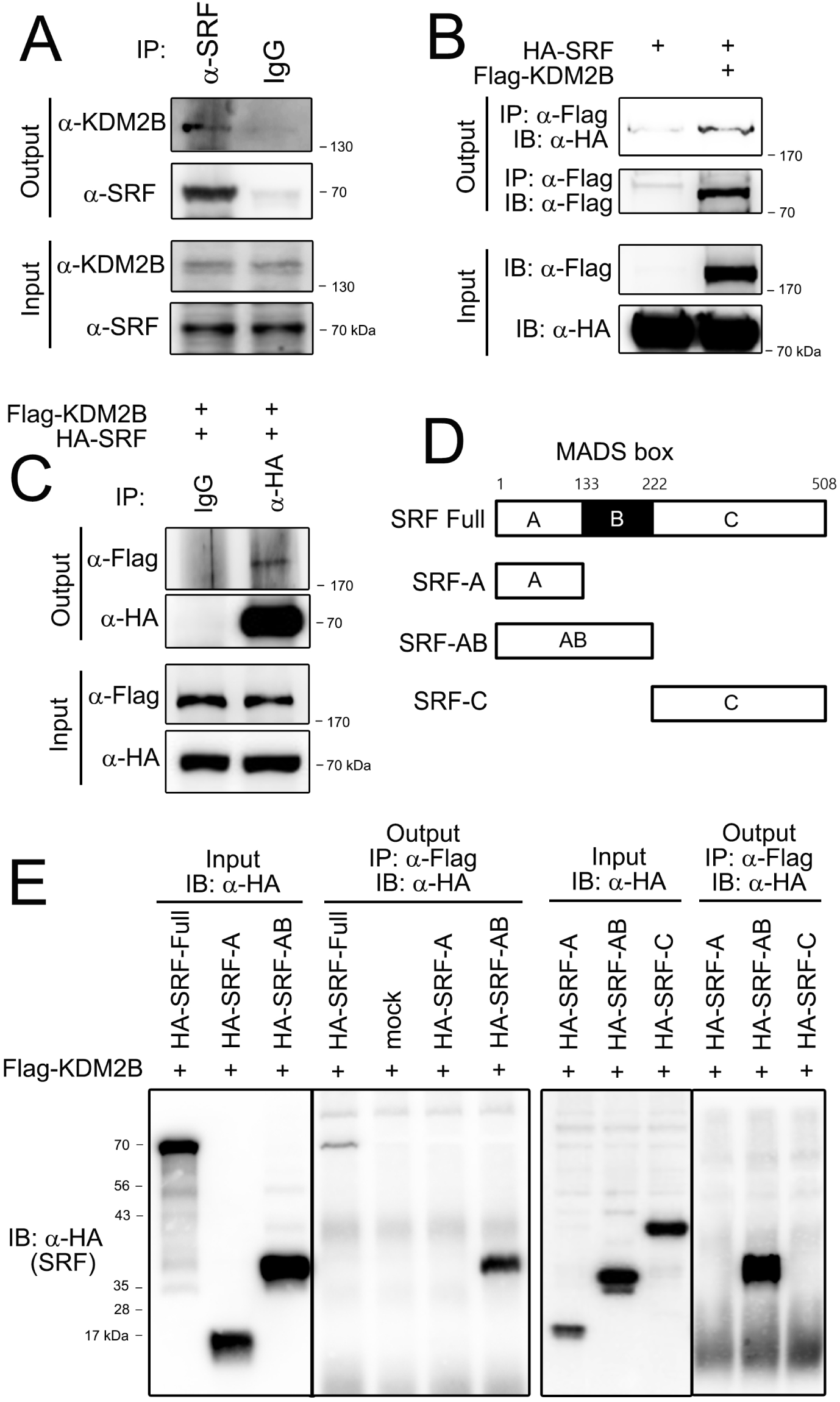
KDM2B directly interacts with SRF through its binding to the MADS box. A. Endogenous SRF successfully recruits endogenous KDM2B in undifferentiated C2C12 cells. B. Flag-tagged KDM2B physically bound to HA-tagged SRF. *pCMV-3xFlag-KDM2B* or *pCGN-HA-SRF* was transfected into 293T cells and immunoprecipitation analysis was performed. C. Inverse immunoprecipitation. HA-tagged SRF pulled down Flag-tagged KDM2B in 293T cells. D. Structure of truncated mutant proteins of SRF. The MADS box consists of the 133-222 amino acid region (black boxed region). E. The MADS box interacts with KDM2B. Left two gels: SRF-full length, SRF-A, and SRF-AB constructs were used for domain-mapping analysis. Immunoprecipitation was performed by anti-Flag (KDM2B) antibody. Right two gels: SRF-A, SRF-AB, and SRF-C constructs were used for immunoprecipitation. Note that only SRF-full length and SRF-AB are recruited by KDM2B.

To determine which domain is responsible for the interaction with KDM2B, we performed domain mapping analysis by using truncated mutant proteins of SRF as shown in Fig 4D. We first examined the expression of SRF-A and SRF-AB truncated mutants (leftmost panel of Fig 4E). The interaction with KDM2B was observed in SRF-AB (4^th^ lane in second panel of Fig 4E) as well as SRF full length (1^st^ lane in the second panel), whereas the interaction was not observed in SRF-A (3^rd^ lane in second panel). We also examined whether SRF-C could interact with KDM2B. All the truncated mutants (SRF-A, SRF-AB, and SRF-C) were well expressed (3^rd^ panel from the left in Fig 4E). However, only SRF-AB could interact with KDM2B (2^nd^ lane in 4^th^ panel from the left in Fig 4E). These results suggested that the MADS box is responsible for the interaction with KDM2B.

We also tested whether KDM2B could interact with MyoD or Pax7, both of which mainly function in the early phase during skeletal myogenesis (Chen *et al*, 2019; Olguin & Pisconti, 2012). Exogenous proteins of MyoD (Fig EV5A) and Pax7 (Fig EV5B) directly interacted with KDM2B.

### SRF MADS box domain is methylated during myocyte differentiation

What would be the function of KDM2B in skeletal muscle differentiation? Would methylation of muscle-specific transcription factors be involved? Does KDM2B demethylate these factors by direct association? To answer these questions, we first checked whether myogenic transcription factors are methylated or demethylated during the differentiation of myoblasts using immunoprecipitation-based methylation analysis. C2C12 cellular lysates were obtained from either GM or DM (2 and 4 days) and subjected to immunoprecipitation with anti-methyl-lysine antibody and the methylated SRF was checked by immunoblot with anti-SRF antibody. DM increased the methylation of SRF (Fig 5A). Next, we wanted to know which SRF domain is responsible for the methylation. To address this, we used truncated SRF mutants. Only SRF-AB was methylated in the DM condition, which suggests that the SRF MADS box is methylated during exposure to DM (Fig 5B).

**Figure 5.**
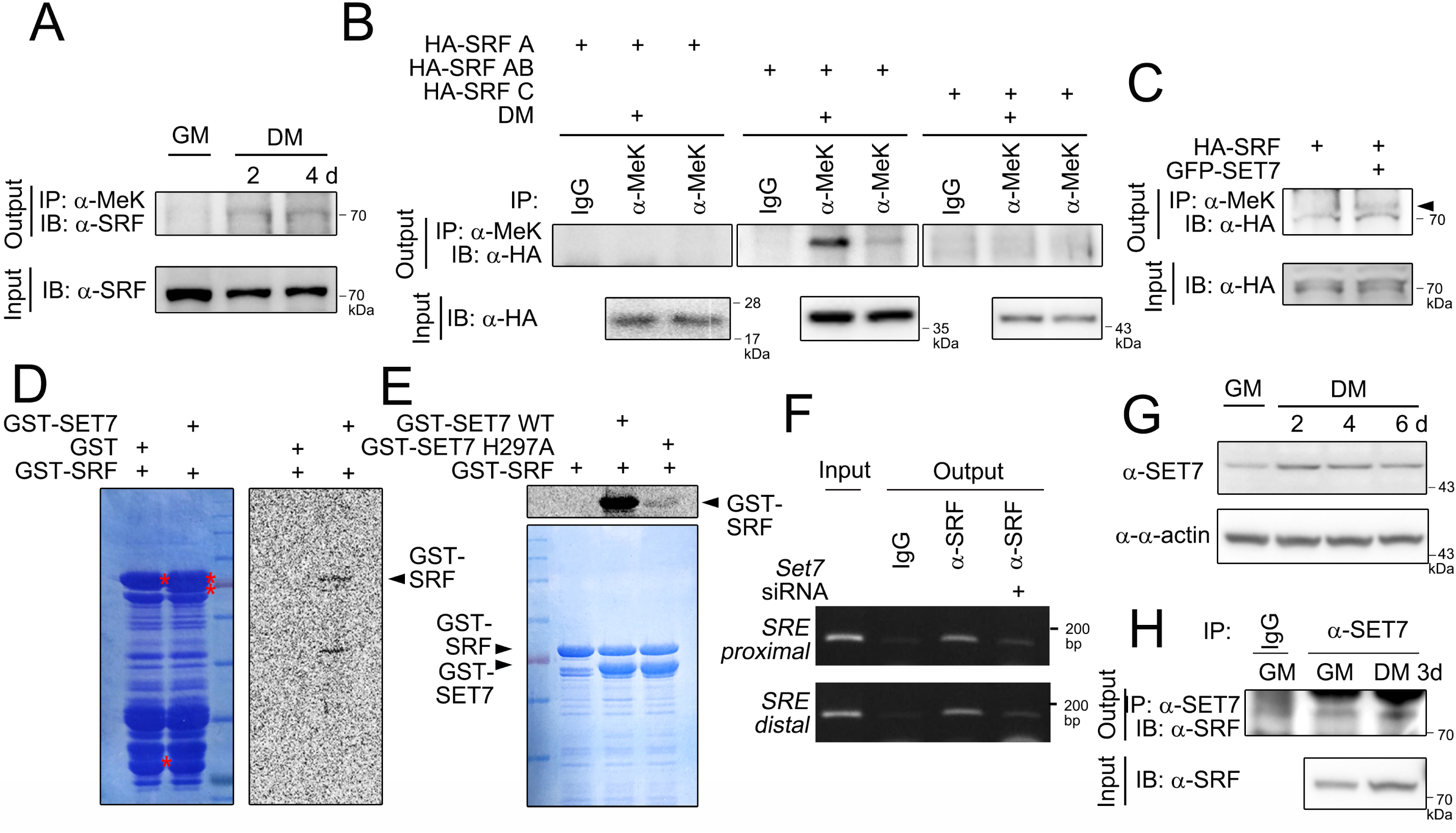
Myoblast differentiation induces SET7-dependent methylation of SRF MADS box. A. Immunoprecipitation-based *in vivo* cellular methylation assay using anti-methyl lysine (α-MeK) antibody revealed SRF is methylated during C2C12 differentiation. Note that methylation is gradually increased with the days of differentiation. B. Truncated mutants of *SRF* (*SRF-A, SRF-AB*, and *SRF-C*) were transfected into C2C12 cells and then the cells underwent serum deprivation. SRF MADS box region is methylated, as determined with α-MeK immunoprecipitation-based assay. C. Transfection of *SET7* induces methylation of SRF in C2C12 cells. D. *In vitro* methyltransferase assay in complete cell-free condition showing that chimeric protein GST-SET7 induces methylation of GST-SRF. Left: Coomassie blue staining to check the \expression of GST proteins. Right: Autoradiograph. E. Enzymatic-activity-dead SET7 (SET7 H297A) fails to induce methylation of SRF. F. *Set7* siRNA reduced binding of SRF to the proximal *SRE* in the *Acta1* gene promoter. ChIP analysis. G. Expression of SET7 is increased during C2C12 differentiation. H. The association between endogenous SET7 and SRF is enhanced in DM.

A previous report by Tuano et al.(Tuano *et al*., 2016) suggested that SET7 acts as a SRF methyltransferase, although it did not appear to have any role in methylation of SRF in myogenesis. Another research group (Tao *et al*., 2011) also showed that SET7 is required for skeletal myogenesis, and that group suggested that SET7-induced methylation of histone H3K9 is required for normal skeletal muscle development (Tao *et al*., 2011). However, the possibility of involvement of a non-histone target of SET7 in the mechanism of action was not demonstrated. In the present work, we examined whether SET7 could methylate SRF. Transfection of *SET7* increased methylation of SRF as determined by immunoprecipitation with anti-methyl lysine antibody (Fig 5C). This was further supported by the results of an *in vitro* methylation assay utilizing S-adenosyl-[methyl-^14^C]-L-methionine [^14^C-SAM]. In contrast to the GST-only condition, a negative control, GST-SET7 induced methylation of GST-SRF (Fig 5D). The SRF-methylation was abolished when enzyme-activity-dead SET7 mutant (SET7 H297A)(Nishioka *et al*, 2002) was used (Fig 5E). ChIP analysis clearly showed that the binding of SRF to the *SRE* is SET7-dependent; transfection of *Set7* siRNA reduced the proximal *SRE* occupancy of SRF in the *Acta1* promoter (Fig 5F and Fig EV6A).

To investigate the involvement of SET7 in myocyte differentiation, we first tested the expression levels of SET7 in C2C12 differentiation. Indeed, the expression of SET7 was increased in the acute phase of myocyte differentiation after exposure to DM for 2 and 4 days (Fig 5G), which was further supported by examination of the mRNA levels (Fig EV6B). The association between SET7 and SRF was enhanced in the DM compared with the GM condition (Fig 5H), which would be caused by the increase in the amount of SET7 and SRF in the DM condition (lower panel input). Transfection of *SET7* further increased SRF-induced transactivation of *Acta1* (Fig EV6C). These results suggested that SET7 is an SRF-methyltransferase and that the methylation of SRF induces transactivation of SRF-mediated skeletal muscle genes in the early phase in myogenesis.

### SRF methylation of lysine residues is induced by SET7 or differentiation media

To investigate the biological role of SRF methylation, we needed to determine which lysine residue is responsible for the SET7-mediated methylation and whether the demethylation is caused by KDM2B. Thus, as shown in the upper table in Fig 6A, we generated 5 synthetic peptides spanning the whole MADS box, which we previously proved as a methylation domain (Fig 5B), and performed an *in vitro* methyltransferase assay. When GST-SET7 was co-incubated, peptide #2, spanning amino acids 153∼167 and which contains K154, K163, and K165, was highly methylated (Fig 6B). To confirm whether the methylation of peptide #2 is dependent on SET methyltransferase activity, we utilized GST-SET7 H297A. SET H297A failed to induce methylation of peptide #2 (Fig EV6D).

**Figure 6.**
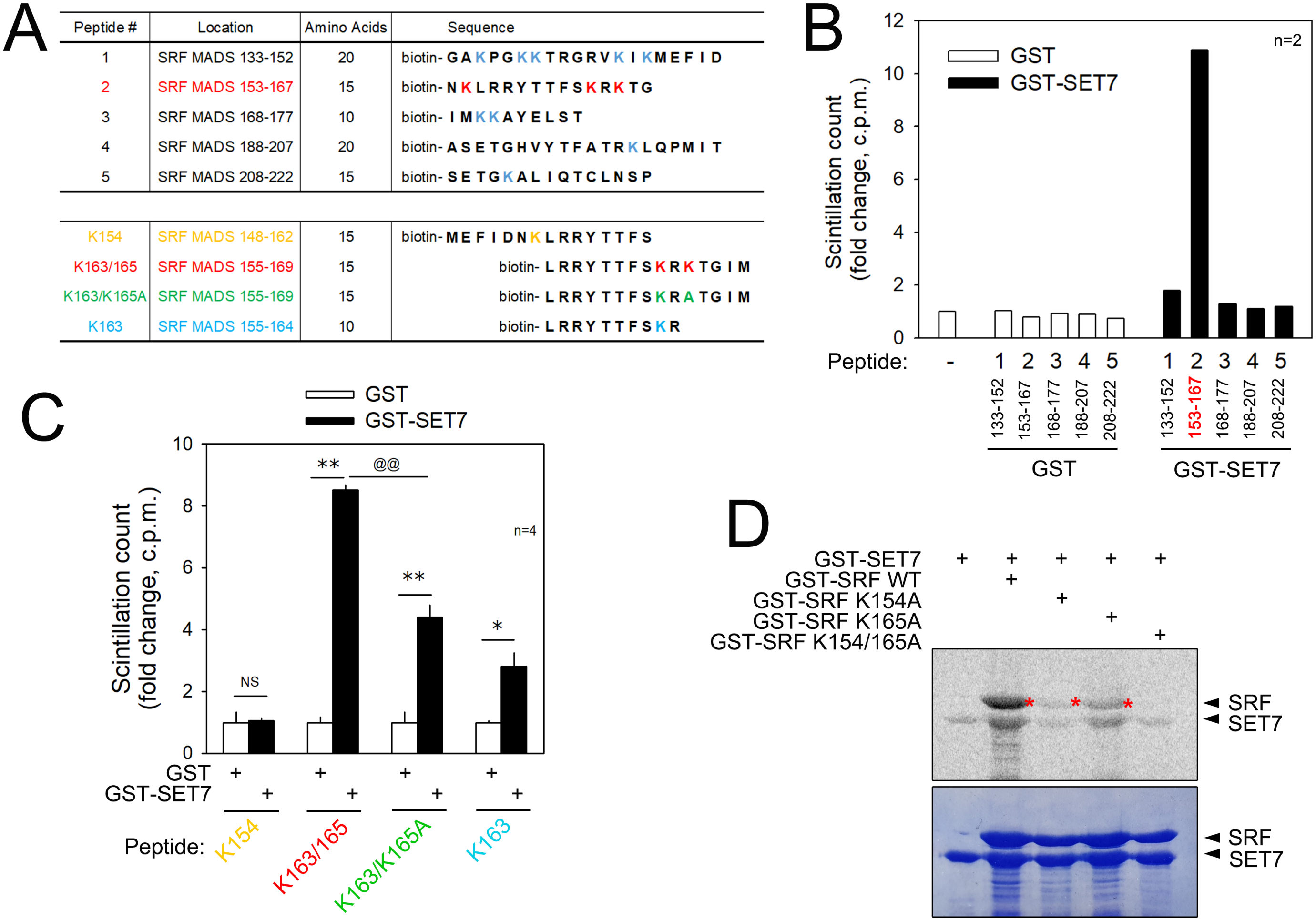
SRF lysine residues that are methylated by SET7. A. Synthetic peptides used in the current study to determine methylated lysine residue. B. Only peptide #2 spanning amino acids 153∼167 in the MADS box of SRF is methylated by GST-SET7. Synthetic peptides were used as substrates and incubated with GST-SET7 for *in vitro* methyltransferase assay. C. To further specify the lysine, K154-, K165-, and K165A-bearing synthetic peptides were generated and used for *in vitro* methyltransferase assay. K154 does not show any methylation, whereas K165 has a strong methylation signal. Note that K165A halved the methylation, which suggests additional methylation other than by K165. D. GST-SRF WT, GST-SRF K154A, GST-SRF K165A, and GST-SRF K154/165A were used for *in vitro* methyltransferase assay. As for the methylation enzyme, GST-SET7 was supplemented. Note that both K154 and K165 are methylated by SET7. ** and @@, p < 0.01.

We next generated two peptides: K154 and K163/165. K154 spans amino acids 148-162, whereas K163/165 contains 155-169 (Fig 6A lower table). In contrast with that of K154 (1^st^ group of bars), the methylation of the peptide spanning K163/165 was significantly increased (2^nd^ group of bars in Fig 6C). The K163/165 peptide spanning 155∼169 contains two lysines. To specify which lysine residue is methylated, we first generated K165-dead mutant by substituting K165 with alanine: K163/K165A (Fig 6A lower table). Interestingly, the methylation of K163/K165A was significantly lower than that of the K163/165 peptide (3^rd^ group of bars in Fig 6C). It is noteworthy that K165A still showed significant methylation; the incomplete elimination of methylation might be caused by SET7-dependent methylation of K163. Again, we generated K163 peptide spanning 155∼164, where only one lysine of K163 was included. K163 was sufficiently methylated by addition of GST-SET7 (4^th^ group of bars in Fig 6C).

We also performed an *in vitro* methyltransferase assay using GST-fusion proteins of SRF. GST-SET7 successfully induced methylation GST-SRF wild type (WT) under *in vitro* conditions (2^nd^ lane of upper gel in Fig 6D and 2^nd^ lane of right gel in Fig EV6E). We also generated GST proteins of SRF K154A, K165A, and K154/165A. Substitution of K154 abolished the methylation of SRF (3^rd^ lane in Fig 6D). The methylation was also significantly reduced in GST-SRF K165A (4^th^ lane in Fig 6D and 3^rd^ lane in Fig EV6E) and in GST-SRF K154/165A (5^th^ lane in Fig 6D). These *in vitro* methylation results suggested that SET7 might induce methylation of K154, K163, and K165. Thus, we further tried to narrow down the specific lysine by examining the biological functions of those residues in relation to myogenesis.

### SRF K165 methylation is critical for SRF transactivation and muscle cell differentiation

In previous studies, we found that SET7 induces methylation of the MADS box domain of SRF and that K154, K163, and/or K165 might be the SET7-dependent methylation targets. It is noteworthy, however, that not all methylations were biologically functional. Thus, we tried to further narrow down the specific methylation residues that are biologically active. To do this, we generated diverse mutants by site-directed mutagenesis in which lysine residues were replaced by alanine residues (Fig EV7A). SRF-induced transactivation of the *Acta1* promoter was completely abolished when *SRF K165A* was transfected (Fig 7A). Interestingly, transactivation was not altered when other lysine residues in the MADS box were mutated, which suggests that K165 is a key lysine in the maintenance of the transcriptional activity of SRF. In addition to basal *Acta1* promoter activity, SRF K165A failed to further increase *Acta1* promoter activity as the SRF wild type did (Fig 7B). The result was similar when a *Myogenin-luciferase* construct was used (Fig 7C). The abolishment of transcriptional activation by SRF K165A was caused by its failure to bind to the *SRE* in the *Acta1* promoter; ChIP analysis showed that *SRE* occupancy by SRF in the *Acta1* promoter was attenuated when *SRF K165A* was transfected (Fig 7D). A representative gel picture is shown in Fig EV7B. Using the gel shift assay, we further studied whether SRF K165 methylation affected the binding of SRF to the distal *SRE* (Fig 7E). Nuclear extract of *SRF K165A*-transfected C2C12 cells failed to form an SRF-SRE complex (4^th^ lane), compared with that of *SRF WT*-transfected cells (3^rd^ lane). Likewise, complex formation between SRF K165A and the proximal *SRE* was significantly reduced (Fig EV7C).

**Figure 7.**
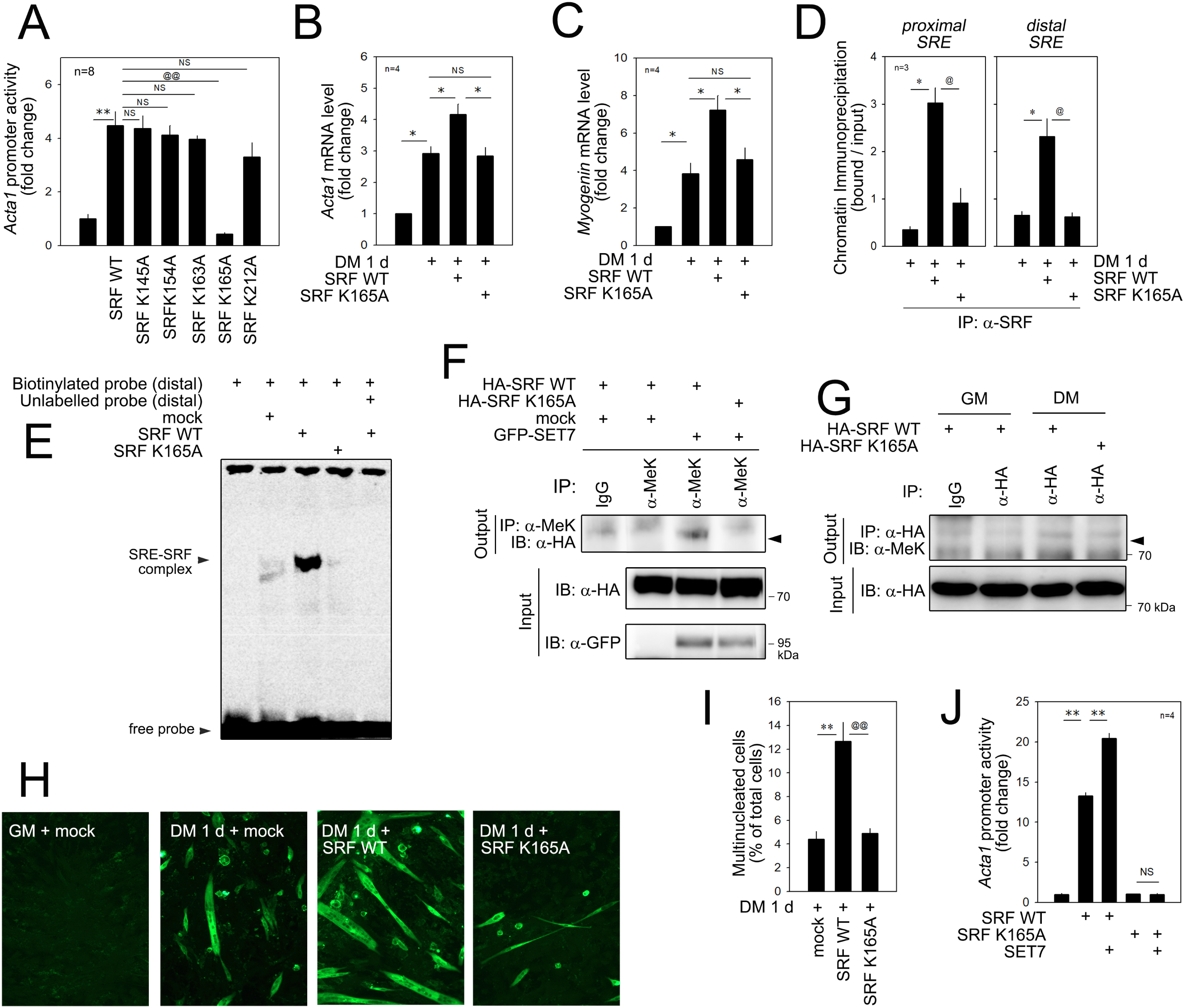
SRF K165 is biologically active methylation site for muscle differentiation. A. Promoter analysis using *Acta1* promoter revealed that only SRF K165A lost its basal transcriptional activity among those tested. B. DM induces *Acta1* mRNA level, which is further potentiated by transfection of *SRF WT*. However, this enhancement is not observed when *SRF K165A* is transfected. C. *Myogenin* mRNA level. D. ChIP analysis showing that SRF K165A loses its binding activity to two *SRE*s (CArG-proximal and CArG-distal) in the *Acta1* promoter. To pull down either SRF WT or SRF K165A, anti-HA antibody was used for ChIP analysis. E. Gel shift assay showing that SRF K165A loses its binding affinity to the distal *SRE*. F. Immunoprecipitation-based *in vivo* cellular methylation assay further showing that the transfection of *SET7* induces methylation of SRF WT, whereas it fails to methylate SRF K165A. G. DM induces methylation of SRF WT, whereas it fails to methylate SRF K165A. H. Elongation and tube formation induced by treatment with DM for 1 d is enhanced by transfection of *SRF WT*, whereas this enhancement is not seen by SRF K165A. Immunocytochemical analysis with anti-MHC antibody. I. Multinucleated cell count. J. Transfection of *SET7* enhances SRF WT-induced transactivation of *Acta1* promoter, but fails to activate for SRF K165A. * and @, p < 0.05; ** and @@, p < 0.01; NS, not significant.

Methylation of SRF was examined by *in vivo* cellular studies; transfection of *SET7* induced methylation of SRF WT (3^rd^ lane of upper gel in Fig 7F). It is noteworthy that SET7 failed to induce methylation in SRF K165A (4^th^ lane). We next examined whether DM also methylated SRF in the absence of forced expression of SET7; treatment of C2C12 cells with DM increased the methylation of SRF WT (2^nd^ vs 3^rd^ lane of upper gel of Fig 7G). However, DM did not induce methylation when SRF K165A (4^th^ lane) was used. These results suggested that at least SRF K165 would be one of the methylation residues in C2C12 differentiation.

The myogenic activity of SRF K165A was examined. Treatment with DM for 1 day induced multinucleation and elongation of C2C12 myoblast cells, which were then enhanced by transfection of *SRF wild type*. However, K165A failed to enhance multinucleation and elongation (Fig 7H). Quantification of multinucleated cells further showed that the substitution of SRF K165 with alanine abolished the SRF-induced increase in multinucleation (Fig 7I). Transfection of *SET7* further increased the SRF-mediated transactivation of *Acta1* promoter (2^nd^ vs 3^rd^ lane in Fig 7J). However, no further activation was observed when *SRF K165A* was transfected (4^th^ vs 5^th^ lane), which suggests that SET7-mediated myogenic gene activation is dependent on SRF K165.

We checked whether substitution of K165 with alanine reduces the binding of SRF to either MyoD or SET7. In our experimental model, the association with those proteins was not altered (Fig EV7D and EV7E).

### KDM2B demethylates SRF K165

Next, we checked whether KDM2B demethylates SRF. The *in vitro* methylation assay showed that the addition of GST-KDM2B attenuated GST-SET-induced methylation of GST-SRF (Fig EV8A) in a dose-dependent fashion (Fig 8A). The *in vitro* methylation assay with synthetic peptide showed that KDM2B demethylates SRF K165; KDM2B reduced SET7-induced methylation when K165 peptide (Fig 8B) as well as peptide #2 spanning 153∼167 amino acids of SRF (Fig EV8B) were used as substrates. In contrast, with the intact K165 peptide whose methylation was significantly reduced by KDM2B (1^st^ vs. 2^nd^ bar), no further reduction of methylation was observed when peptide K165A was used as a substrate (3^rd^ vs. 4^th^ bar, Fig EV8C). Treatment with DM increased the methylation of endogenous SRF in C2C12 cells, as determined by an *in vivo* immunoprecipitation-based methylation assay (3^rd^ lane in Fig 8C). The increase in methylation was blunted when *KDM2B* was transfected into the C2C12 cells (4^th^ lane in Fig 8C).

**Figure 8.**
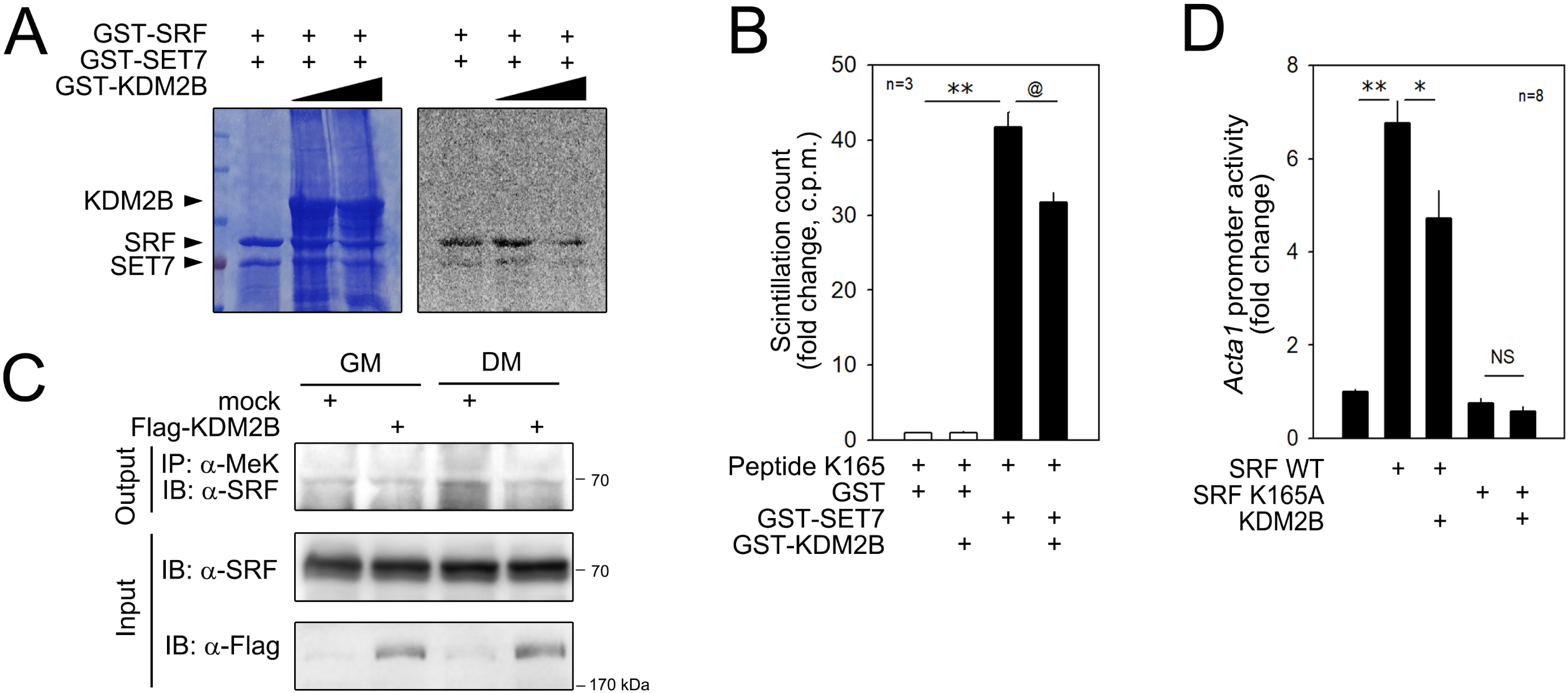
KDM2B demethylates SRF WT to mimic SRF K165A. A. GST-KDM2B reduces the methylation of SRF in a dose-dependent fashion. *in vitro* methylase assay. B. *in vitro* methylase assay with synthetic peptide. GST-SET7 induced methylation of peptide K165, which is partially reduced by addition of GST-KDM2B. C. Immunoprecipitation-based *in vivo* methylation assay. DM-induced methylation of SRF is reduced by transfection of *KDM2B* in C2C12 cells. D. Transfection of *KDM2B* reduces SRF WT-induced transactivation of *Acta1* promoter, whereas it fails to activate for SRF K165A. **p < 0.01, @p < 0.05, @@p < 0.01.

We next questioned whether the KDM2B-mediated removal of SET7-induced methylation resulted in transcriptional regulation in the *in vivo* cellular conditions. As shown in Fig 7I, SET7 further enhanced SRF-induced transactivation of the *Acta1* promoter. This enhancement, however, was abolished by co-transfection of *KDM2B* in a dose-dependent fashion (Fig EV8D).

To learn whether SRF K165A mimics demethylated SRF, we first assumed that KDM2B would not further inhibit the transcriptional activity of SRF K165A if K165 was the only target of KDM2B in transcriptional regulation. As shown in Fig EV2A to EV2D, SRF WT-induced transactivation of the *Acta1* promoter was blunted by transfection of *KDM2B* in C2C12 cells (2^nd^ and 3^rd^ bars, Fig 8D). SRF K165A failed to transactivate the *Acta1* promoter as shown in Fig. 7a (5^th^ bar). Importantly, *KDM2B* transfection did not further reduce SRF K165A activity, as it did SRF WT activity (4^th^ and 5^th^ bar, Fig 8D). Together with the failure of Set7-mediated potentiation of SRF K165A-transactivation (Fig 7I), these results suggested that the intact SRF K165, at least in part, mediated the effect of SET7 and KDM2B.

In the present work, we showed that both MyoD and Pax7 could interact with KDM2B (Fig EV5). In contrast to SRF, however, neither transcription factor was ‘demethylated’ by KDM2B; rather, the methylation of these two transcription factors seemed to be increased (Fig EV8E and EV8F). Although the reason for the ‘hypermethylation’ is not clear, these results led us to rule out investigation of these muscle-specific transcription factors as targets of KDM2B.

### SET7 inhibitors mimic KDM2B

We wanted to find out if pharmacologic inhibition of SET7 might attenuate skeletal muscle differentiation like KDM2B did. Both sinefungin (sine)(Sasaki *et al*, 2016; Tamura *et al*., 2018) and (R)-8-fluoro-N-(1-oxo-1-(pyrrolidin-1-yl)-3-(3-(trifluoromethyl)phenyl)propan-2-yl)-1,2,3,4-tetrahydroisoquinoline-6-sulfonamide hydrochloride [(R)-PFI-2, or R-PFI](Barsyte-Lovejoy *et al*, 2014) are known to inhibit SET7. Treatment with sine did not affect C2C12 cell survival. However, R-PFI induced proliferation (Fig EV9A). Sine dose-dependently inhibited *Acta1* promoter activity (Fig 9A). Treatment of C2C12 with sine attenuated the binding of SRF to either the proximal or the distal *SRE* (Fig 9B and 9C). The sine-induced reduction of SRF-dependent transactivation resulted in the decreased expression of mRNAs of myogenic genes (Fig 9D) and their protein products (Fig 9E). Indeed, sine reduced the expression of MHC (Fig 9F) and the number of multinucleated C2C12 cells (Fig 9G).

**Figure 9.**
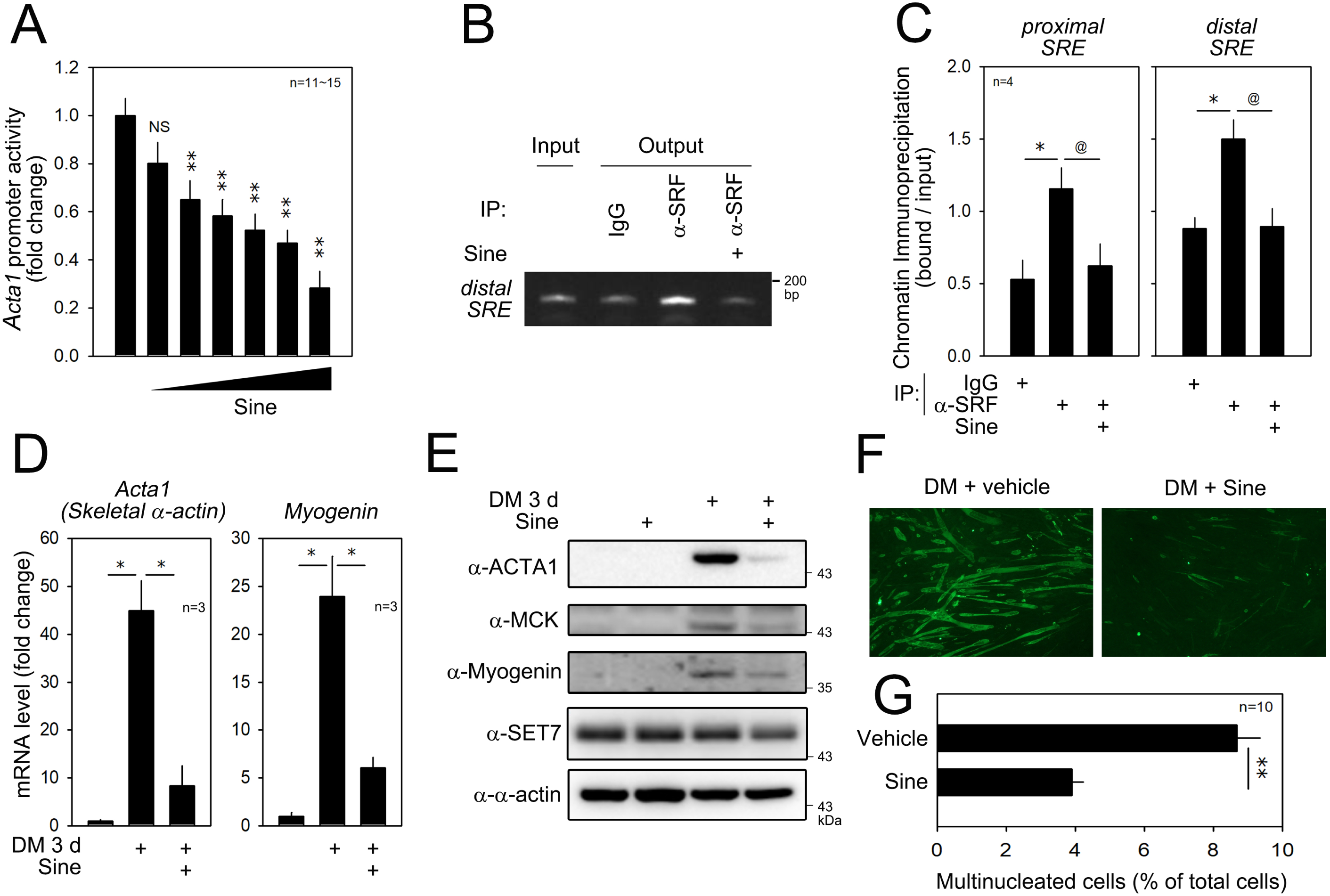
Sinefungin (sine), an SET7 inhibitor, attenuates myoblast differentiation by transcriptional repression of SRF-dependent genes. A. Sine inhibits *Acta1* promoter activity in a dose-dependent fashion. B. Representative gel picture for ChIP. Sine reduces the binding of SRF to the distal *SRE* in the *Acta1* gene promoter. C. Quantification results of ChIP analysis. D. Quantitative RT-PCR results showing that sine abolishes the increases of *Acta1* and *Myogenin* mRNAs induced by DM for 3 d. E. Sine blocks the increases in the amounts of ACTA1, MCK, and Myogenin induced by DM for 3 d. F. Immunofluorescent images of *α*-MHC. G. Multinucleated cell count. *p < 0.05, **p < 0.01.

R-PFI, an alternate SET7 inhibitor, also inhibited the activity of the *Acta1* promoter (Fig EV9B) as well as the binding capacity of SRF to the *SRE* (Fig EV9C and EV9D). mRNA levels and subsequent protein levels of both *Acta1* and *Myogenin* were reduced by treatment with R-PFI (Fig EV10A and EV10B). R-PFI also inhibited the expression of MHC proteins (Fig EV10C) and led to a decrease in multinucleated cells (Fig EV10D). These results suggested that Set7 inhibitors also block myogenic differentiation in an SRF-mediated transactivation-dependent manner.

## Discussion

In the present work, we have shown a novel function of the methylation of SRF in myogenic differentiation. We observed that KDM2B inhibits myogenic differentiation by repressing SRF-dependent transcriptional activity. The transcriptional repression was not caused by the demethylation of histones. Instead, we found that SRF, a non-histone target, is methylated during myogenic differentiation in a SET7-dependent fashion and that KDM2B-induced demethylation contributes to transcriptional repression by inducing the detachment of SRF from the *SRE*. SET7, not KDM2B, was shown to methylate SRF K165. Thus, the SET7/KDM2B-mediated balance of K165 methylation is important in regulating the transcriptional activity of SRF by adjusting its nuclear localization (Fig 10).

**Figure 10.**
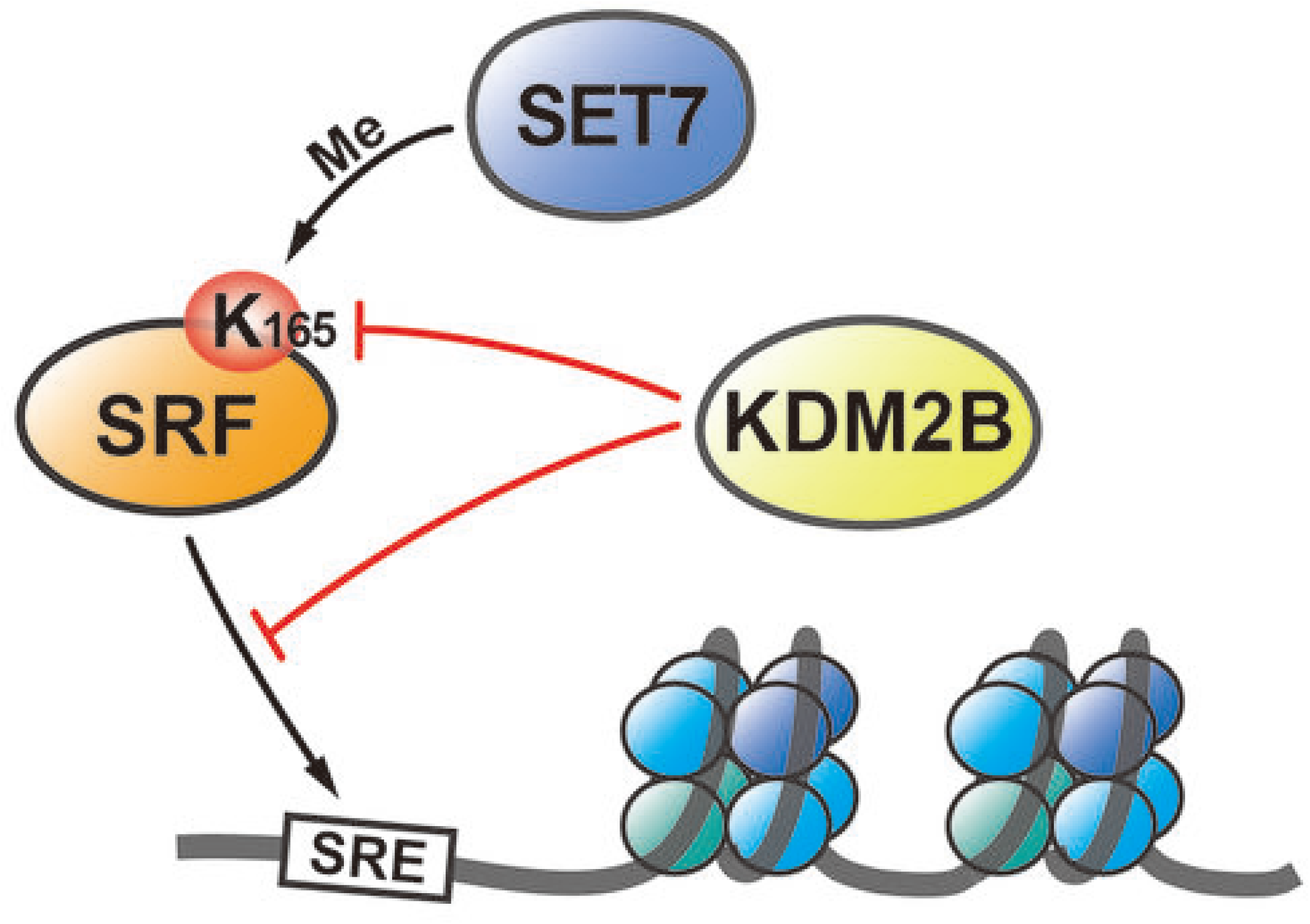
Diagram. Working hypothesis. KDM2B inhibits skeletal muscle differentiation by inhibition of SRF-induced transactivation of skeletal muscle genes. KDM2B directly binds to SRF to demethylate K165, which results in detachment of SRF from the *SRE* in skeletal muscle-specific genes. SET7 counteracts KDM2B by methylation of SRF K165.

Muscle differentiation is a well-orchestrated sequence from mesodermal progenitor cells to muscle fibers (Li *et al*, 2005). Under the influence of many regulators, myoblasts form fiber-shaped myotubes by fusing with neighboring myocytes to make mature multinucleated myocytes (Chal & Pourquie, 2017). Diverse epigenetic regulators and chromatin-modifying enzymes are known to regulate these processes. Importantly, these epigenetic modifiers finely regulate muscle differentiation not only by their histone modification, but also by PTMs of muscle-specific transcription factors. In particular, several recent reports have indicated that histone methyltransferases and demethylases also target non-histone substrates in myogenesis. For example, SET7 mediates histone H3K9 methylation (Tao *et al*., 2011), whereas lysine methyltransferase G9a methylates the key myogenic transcription factor MyoD (Ling *et al*, 2012). Moreover, G9a methylates K267 of MEF2D and represses its transcriptional activity, while LSD1 demethylates it to induce myogenic differentiation (Choi *et al*, 2014; Tosic *et al*, 2018). KDM4A, an alternate histone demethylase, is required for skeletal muscle differentiation(Verrier *et al*, 2011) and regulates myogenesis in C2C12 cells by interacting with MyoD (Choi *et al*, 2015). In the present study, we found that KDM2B inhibits myogenesis. However, it is noteworthy that KDM2B inhibits myogenic differentiation in a histone-methylation-independent manner.

SRF is also susceptible to diverse PTMs such as phosphorylation and glycosylation (Janknecht *et al*, 1992; Reason *et al*, 1992). These modifications are related to homeostasis and pathologic conditions (Blaker *et al*, 2009; Iyer *et al*, 2006). Phosphorylation of S162 in the SRF MADS box regulates proliferation and myogenic gene programs. Furthermore, PKCα phosphorylates SRF S162 (Iyer *et al*., 2006). SRF can also be O-GlcNAc-modified (Schroter *et al*, 1990). Using recombinant SRF overexpressed in baculovirus, Marais et al.(Marais *et al*, 1992) identified S283 as a major site of GlcNAc modification. S316 was also found to be glycosylated, and one of either S307 or S309 sites appeared to be modified. SRF is modified with SUMO-1 at K147 *in vivo* and *in vitro* (Matsuzaki *et al*, 2003).

Although it had been predicted that SET7 might methylate SRF,(Tuano *et al*., 2016) there had been no experimental validation of this. In addition, the function of SET7-mediated methylation of SRF had never been documented. In this study, however, we clearly showed that SET7 works as an SRF-methyltransferase. SET7-induced histone methylation has been emphasized in muscle biology (Tao *et al*., 2011). It was observed that SET7 induces monomethylation of histone (H3K4me1), which results in the recruitment of SRF at the promoter. Here, we suggest that SET7 counteracts KDM2B in regulating the methylation of SRF K165 and thereby modulates occupancy of the *SRE* for transcriptional activation. Although it is known that KDM2B mediates demethylation of both histone and non-histone substrates, the biological significance of non-histone demethylation, especially in skeletal muscle differentiation, was not previously known. Our data provide unique insights into the role of demethylation of SRF, a non-histone substrate, in the control of myogenic differentiation. Moreover, our data indicate that SRF methylation, without alteration in histone methylation, plays a significant role in KDM2B-mediated inhibition of myogenesis.

## Materials and Methods

### Antibodies and reagents

Antibodies against SRF (sc-335), MyoD (sc-304), Set7/9 (sc-390823), MCK (sc-15164), and GAPDH (sc-166574) were obtained from Santa Cruz Biotechnology (Santa Cruz, CA, USA). Antibodies against Flag (F1804), HA (H9658), GFP (G1544), and actin (A2066) were purchased from Sigma-Aldrich (St. Louis, MO, USA). Antibodies against histone H3 (tri-methyl K4, ab8580), histone H3 (di-methyl K36, ab9049), pan-methyl lysine (ab7315), and lamin B1 (ab16048) were from Abcam (Cambridge, UK). Anti-KDM2B/JHDM1B antibody (09-864) was from EMD Millipore Corp (Billerica, MA, USA). Anti-MHC antibody (MF-20) was from DSHB (Iowa City, IA, USA). Anti-V5 antibody (46-0705) was from Invitrogen (Carlsbad, CA, USA). SRF-derived peptides (1∼5, K154, K163, K165, K165A) were synthesized by Peptron (Daejeon, South Korea). Sinefungin and (R)-PFI-2 were purchased from Sigma-Aldrich (St. Louis, MO, USA).

### Plasmid constructs and transfection

*pCGN-SRF-HA* and *pcDNA3-HA-Pax7* were kindly provided by Prof. Jonathan A. Epstein (University of Pennsylvania, Philadelphia, PA, USA). *pCMV-3xFlag-KDM2B* was described previously.^16^ *pHM6-HA-MyoD* was obtained from Prof. Young Kyu Ko (College of Life Sciences and Biotechnology, Korea University, Seoul, South Korea). Mutants of SRF were constructed by site-directed mutagenesis based on *pCGN-SRF-HA* wild-type (CosmoGeneTech, Seoul, South Korea). Three SRF truncated mutants of SRF A domain-containing region (amino acids 1 to 133), SRF B domain-containing region (amino acids 133 to 222), and SRF C domain-containing region (amino acids 222 to 508) are shown in Fig. 4D. *pGL3-basic*-*myogenin-luciferase* and *pGL3*-*Mck-luciferase* were kind gifts from Prof. Da-Zhi Wang (University of North Carolina, NC). For the *Acta1* promoter-reporter assay, −450 to +26 base pairs from the transcription start site were amplified from mouse genomic DNA and subcloned into *pGL3* basic vector.

C2C12 cells were transfected using lipofectamine and Plus reagent (Invitrogen, Carlsbad, CA, USA) according to the manufacturer’s directions. Each recombinant expression vector was transiently transfected into HEK293T cells with PEI reagents according to the manufacturer’s instructions. The siRNA against KDM2B was obtained from Bioneer (Daejeon, South Korea). Cells were transfected with siRNA (50 nM) using lipofectamine RNAi MAX (Invitrogen, Carlsbad, CA, USA) according to the manufacturer’s instructions.

### Cell culture and subcellular fractionation

C2C12 cell lines were maintained in Dulbecco’s modified Eagle’s medium (DMEM) supplemented with 15% fetal bovine serum (FBS) and antibiotics. DMEM with 2% horse serum was used as the differentiation medium. HEK293T cells were maintained in DMEM supplemented with 10% FBS and antibiotics, as previously described. To separate nuclear proteins, NE-PER nuclear and cytoplasmic extraction reagents (Thermo Fisher Scientific, Waltham, MA, USA, #78833) were used.

### RNA isolation and real-time quantitative PCR (q-PCR)

Total RNA was isolated from cells using NucleoSpin kits (Macherey-Nagel, Bethlehem, PA, USA). One microgram of total RNA was used for q-PCR analysis. cDNA was synthesized using ReverTra Ace cDNA synthesis kits (TOYOBO, Nipro, Osaka, Japan) and 1 μL of the cDNA synthesis reaction mixture was used with the Platinum SYBR Green kit from Qiagen (Hilden, Germany). q-PCR was performed using a Rotor-Gene Q Real-time PCR machine (Qiagen, Hilden, Germany). The oligonucleotides used in PCR were Kdm2b forward, 5’-AGCAGCTAAAACCTGGCAAA-3’, and reverse, 5’-GTGAGCTGGAACGTGACTGA-3’; Acta1 forward, 5’-GACCTCACTGACTACCTGATGAAA-3’, and reverse, 5’-CAGACTCCATACCGATAAAGGAAG-3’; myogenin forward, 5’-AGTACATTGAGCGCCTACAG-3’, and reverse, 5’-ACCCACCCTGACAGACAATC-3’; Mck forward, 5’-AGCAGCTCATTGATGACCAC-3’, and reverse, 5’-TCAAACTTGGGGTGCTTGCT-3’; beta-actin forward, 5’-CACGATGGAGGGGCCGGACTCATC-3’, and reverse, 5’-TAAAGACCTCTATGCCAACACAGT-3’.

### Immunoblot analysis

Cell lysates were obtained from cells and tissues using RIPA buffer (R2002, Biosesang, Seongnam, South Korea) or 0.5% NP lysis buffer supplemented with protease inhibitors. Approximately 20–30 μg total lysates of cell lines or IP-eluted samples were separated by 10% or 12% SDS-PAGE gels and subsequently transferred to PVDF membranes (Millipore, Billerica, MA, USA). Membranes were blocked in 5% skim milk in TBST (0.05% Tween20) buffer for 1 h at room temperature and then incubated with the primary antibodies in 5% milk in TBST buffer overnight at 4 °C. After washing in TBST 3 × for 5 min each, the membranes were incubated with the secondary antibody anti-mouse IgG and anti-rabbit IgG conjugated with horseradish peroxidase (HRP) for 1 h at room temperature. After washing in TBST 3 × for 10 min each, the membranes were incubated with ECL reagent (Millipore, Billerica, MA, USA). Analysis of western blots was conducted using a c300 system (Azure Biosystem, Inc, Dublin, CA, USA). In addition, mouse skeletal muscle developmental western blots (MW-102-d, Zyagen, San Diego, CA, USA) and mouse normal tissue blot II (1562; ProSci, San Diego, CA, USA) were blotted with the same conditions.

### Protein immunoprecipitation (IP)

HEK293T and C2C12 cells were lysed in 0.5% NP40 lysis buffer supplemented with a protease inhibitor. About 1∼2 mg total lysate was incubated with 2 μg antibody or IgG (Santa Cruz Biotechnology, Santa Cruz, CA, USA) coupled to protein A/G agarose beads (Santa Cruz Biotechnology, Santa Cruz, CA, USA) overnight at 4 °C. The beads were washed with ice-cold lysis buffer three times, and the proteins were then eluted from the beads with SDS buffer and subjected to SDS-PAGE.

### Luciferase reporter gene assay

For the luciferase assay, 293T and C2C12 cells were plated on 24-well plates and cultured for 24 h prior to transfection. Cells were transfected with plasmids containing the *pCMV-beta-galactosidase. pCMV-beta-galactosidase* was used to normalize the luciferase assay. Forty-eight hours after transfection, cells were washed with PBS, dissolved in reporter lysis buffer (Promega, Madison, WI, USA), and harvested by scraping. Lysate samples were mixed with luciferase assay reagent and measurements were performed using a luminometer.

### Bacterial expression of GST-fusion proteins

SRF, SRF-K165A, SET7, and KDM2B_1–734_ were cloned into pGEX-4T-1 (GST tag vector) (GE Healthcare, Marlborough, MA, USA). The BL21 strain of *E. coli* was transformed with the target constructs. Transformants of SRF, SRF-K165A, and SET7 were grown in 2X YT medium (16 g/L tryptone, 10 g/L yeast extract, 5 g/L NaCl) until the OD600 was ∼0.6 and induced with 0.02 mM IPTG at 37 °C for 4 h. In the case of transformants of KDM2B_1–734_, induction was using 30°C for 4 h. Bacteria were harvested, and GST-tagged target protein was purified using Glutathione Sepharose 4B (GE Healthcare, Marlborough, MA, USA) according to the manufacturer’s instructions.

### *In vitro* methyltransferase assay

Recombinant GST-SRF protein or synthetic SRF peptides were incubated for 3 h at 30 °C with GST-SET7 in the presence of 100 nCi of S-adenosyl-[methyl-^14^C]-L-methionine [^14^C-SAM] (Perkin Elmer, Waltham, MA, USA) in HMTase assay buffer (50 mM Tris–HCl [pH 8.5], 20 mM KCl, 10 mM MgCl_2_, 10 mM β-mercaptoethanol, and 1.25 M sucrose). Reaction products were separated by SDS-PAGE and analyzed by Phosphorimager (Bio-Rad, Irvine, CA, USA).

### *In vitro* demethylase assay

Recombinant GST-SRF protein or synthetic SRF peptides were incubated for 6 h at 37 °C with GST-KDM2B_1–734_ in demethylation assay buffer (20 mM Tris-HCl, pH 7.3, 150 mM NaCl, 1 mM a-ketoglutarate, 50 mM FeSO_4_, and 2 mM ascorbic acid) following ^14^C-labeling using GST-SET7. Reaction products were separated by SDS-PAGE and analyzed by Phosphorimager.

### Scintillation counting

Synthetic SRF peptides were subjected to an *in vitro* methyltransferase assay using GST-SET7. The ^14^C-labeled peptides were transferred onto p81 filter paper (Millipore) and washed three times with 95% ethanol for 5 min at room temperature. The filters were allowed to air-dry, after which 2 mL of Ultima Gold (Perkin Elmer, Waltham, MA, USA) was added. ^14^C-SAM was then quantified using a scintillation counter.

To measure the remaining radioactivity after the *in vitro* demethylase assay, biotin-conjugated SRF peptides were ^14^C-labeled using GST-SET7, pulled down using streptavidin beads, and then incubated overnight at 37 °C with GST-KDM2B_1–734_ in demethylation assay buffer. The beads were washed, after which 1 mL of Ultima Gold was added, and ^14^C-SAM was quantified using a scintillation counter.

### Chromatin immunoprecipitation assay

Chromatin immunoprecipitation (ChIP) assays were conducted with an EpiQuik Chromatin Immunoprecipitation Kit (Epigentek, Farmingdale, NY, USA) according to the manufacturer’s protocol. Briefly, C2C12 cells were treated with 1% formaldehyde for 10 min to induce cross-links between proteins and DNA, which interacts within intact chromatin. The cells were then sonicated to shear the chromatin fragments to 100 to 500 bp. The sonicated chromatin was immunoprecipitated with anti-SRF, anti-H3K4me3, anti-H3K36me2, and anti-HA, while the negative control was immunoprecipitated with non-immunized IgG. The oligonucleotides used in ChIP PCR were *SRE* proximal forward, 5’-AGTCCTCTCCTTCTTTGGTCAGT-3’, and reverse, 5’-TCCCCTTGCACAGGTTTTTAT-3’; *SRE* distal forward, 5’-GGGCTTATTTTCCATCCCTACC-3’, and reverse, 5’-GTTTGAAAGGTCTCCCCAGTTC-3’.

### Gel shift assay

A nonradioactive LightShift Chemiluminescent EMSA kit (Thermo Fisher Scientific, Waltham, MA, USA, #20148) was used to detect whether SRF binds to the putative SRF binding sites (*SRE*) on the Acta1 proximal promoter. The target oligonucleotide was 3’ end labeled with biotin using a Biotin 3’ End DNA labeling kit (Thermo Fisher Scientific, Waltham, MA, USA, #89818). The oligonucleotides used as probes or competitors in gel shift assays were end-labeled at their 3’ ends with biotin. The sequences were as follows: *SRE* Near: sense, 5’-GACACCCAAATATGGCTTGG-3’; antisense, 5’-CCAAGCCATATTTGGGTGTC-3’ and *SRE* Far: sense, 5’-AGAACCCATAAATGGGGTGC-3’; antisense, 5’-GCACCCCATTTATGGGTTCT-3’. The complementary oligonucleotide pairs were annealed into double strands. Nuclear cell extracts were incubated with biotin-labeled probe in binding buffer (included with EMSA kit) for 30 min at room temperature. An unlabeled oligonucleotide with the same sequence was used as the competitor. Supershift analysis was performed by adding SRF antibody. Loading buffer was added to the reactions, and the binding reaction mixtures were subjected to electrophoresis on 5% PAGE gels in 0.5X TBE buffer at 100 V for 1 h. After electrophoresis, the binding reactions were then transferred onto Biodyne B Nylon membranes (Thermo Fisher Scientific, Waltham, MA, USA, #77016) using 0.5X TBE for 1 h at 60 V. At the end of transfer, the transferred DNA-protein complexes were then cross-linked onto membranes using a hand-held UV lamp equipped with 254-nm bulbs that were set at a distance of approximately 0.5 cm from the membrane for an exposure time of 5-10 min. The blot was visualized by use of a c300 system (Azure Biosystem, Inc, Dublin, CA, USA).

### Immunofluorescence analysis and multinucleated cell counting

C2C12 cells were seeded for fusion or differentiation analysis in triplicate. Cells were fixed in 2% paraformaldehyde for 10 min at room temperature, permeabilized with 0.2% triton for 5 min at room temperature, and blocked with 2–3% normal goat serum for 30 min at room temperature. Primary antibodies for MHC (DSHB, Iowa City, IA, USA) were diluted 1:50 in blocking buffer and incubated at room temperature for 2 h; cells were then incubated with Alexa-conjugated secondary antibody (Molecular Probes, Invitrogen, Carlsbad, CA, USA). After washing, the nuclei were stained with DAPI (p36935, Invitrogen, Carlsbad, CA, USA). The stained cells were analyzed using a fluorescence microscope. Either the total number of nuclei or the number of nuclei within MHC-positive myotubes was counted within 10 individual fields per well. The fusion index was determined by the calculation: Fusion index (%) = (number of nuclei within MHC-stained myotubes/total number of nuclei) ×100. All experiments were performed in triplicate.

### Statistical analysis

Data are represented as means ± SEM. The data were analyzed using either unpaired Student’s *t* test or one-way analysis of variance, which was followed by the Tukey’s honestly significant difference (HSD) multiple-comparison post-hoc test. Statistical analysis was performed with PASW Statistics 21 (SPSS, IBM Company, Chicago, IL). Differences were considered to be significant when p < 0.05.

## Acknowledgements

This research was supported by the National Research Foundation (NRF) of Korea, funded by the Ministry of Education (2018R1A2B3001503), by an NRF grant funded by the Korean government (MSIT) (2017R1D1A1B03031072), and by an NRF grant, Ministry of Science, ICT & Future Planning, Korea (NRF-2019R1A4A2001609 and 2019R1A4A1028534). The authors would like to thank Jennifer Holmes at Medical Editing Services for language editing and careful reading of the manuscript.

## Author contributions

H.J., J.-Y. K., and A.J. performed most experiments. J.-Y.Kang and J.-Y.Kim performed the *in vitro* methyltransferase assay. H.J. and D.-H.K. performed the chromatin immunoprecipitation assays. S.S. and Y.-G.L. performed qRT-PCR and promoter analysis. H.J. and H.-K. M. performed immunoprecipitation. Y.-K.K. performed all bioinformatics analysis. H.J., J.-Y.Kang, J.-Y.Kim, D.-H.K., A.J., Y.-K.K., and S.-B.S. discussed the results and gave critical comments on the manuscript. H.K. designed the research and provided funding. H.J., S.B.S., and H.K. wrote the manuscript.

